# Physiology of the Weight Loss Plateau after Calorie Restriction, GLP-1 Receptor Agonism, and Bariatric Surgery

**DOI:** 10.1101/2023.11.05.565699

**Authors:** Kevin D. Hall

## Abstract

**Objective:** To investigate why different weight loss interventions result in varying durations of weight loss prior to approaching plateaus.

**Methods:** A validated mathematical model of energy balance and body composition dynamics was used to simulate mean weight loss trajectories in response to intensive calorie restriction, semaglutide 2.4 mg, tirzepatide 10 mg, and Roux en-Y gastric bypass (RYGB) surgery interventions. Each intervention was simulated by varying two model parameters affecting energy intake to fit the observed mean weight loss data. One parameter represented the persistent magnitude of the intervention to shift the system from baseline equilibrium and the other parameter represented the strength of the feedback control circuit relating weight loss to increased appetite.

**Results:** RYGB surgery resulted in a persistent intervention magnitude more than 4-fold greater than calorie restriction and about double that of tirzepatide and semaglutide. All interventions except calorie restriction substantially weakened the appetite feedback control circuit resulting in an extended period of weight loss prior to the plateau.

**Conclusions:** These preliminary mathematical modeling results suggest that both GLP-1 receptor agonism and RYGB surgery interventions act to weaken the appetite feedback control circuit regulating body weight and induce greater persistent effects to shift the body weight equilibrium as compared to intensive calorie restriction.

## Introduction

Every obesity intervention eventually results in a body weight plateau after which no further weight loss occurs. The timing of the plateau is a subject of great interest, especially in the context of the recently introduced GLP-1 receptor agonists that exhibit ongoing weight loss without an obvious plateau until well after 12 months (1, 2). Similarly, bariatric surgery often results in a prolonged period of weight loss whereas diet interventions typically exhibit plateaus within 6-12 months. What explains these differing weight trajectories?

Here, I used a validated mathematical model of energy balance and body composition dynamics (3) to simulate the weight loss kinetics within the context of a physiological system that regulates body weight via feedback control of both energy intake and expenditure (4, 5). Specifically, I sought to quantify the typical magnitude of calorie restriction, semaglutide, tirzepatide, and Roux en-Y gastric bypass (RYGB) surgery interventions by simulating published data. Preliminary results suggest that, unlike calorie restriction, semaglutide, tirzepatide, and RYGB weaken the appetite feedback control circuit resulting in an extended period of weight loss prior to the plateau.

## Methods

A previously published and validated mathematical model of human energy metabolism and body composition dynamics (3, 6) was recently modified to represent feedback control of energy intake in response to interventions that induce negative energy balance (4, 5). Since most weight loss interventions in humans mainly affect energy intake, I implemented interventions as follows:

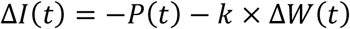

where ΔI(t) is the change in energy intake over time relative to the weight maintenance baseline, P(t) is a parameter representing the overall strength of the intervention, and k is a positive feedback gain parameter relating appetite to weight change, ΔW(t). In the absence of an intervention, P(t) = 0 and k = 95 kcal/d per kg corresponding to the baseline strength of the feedback circuit controlling appetite as previously estimated from modeling weight loss trajectories during sodium glucose co-transporter 2 inhibition (5).

Mathematical model simulations were initialized using mean baseline anthropometric data from published studies on calorie restriction (7), RYGB surgery (8), semaglutide (2), and tirzepatide (1) assuming a mean baseline free-living physical activity level of 1.65 (9). Each intervention was simulated by fitting the parameters P(t) and k such that the modeled body weight time courses matched the observed mean weight trajectories. Body fat was measured by dual energy X-ray absorptiometry in all published intervention studies and mean body fat data was compared to model simulations that assumed nonlinear partitioning between lean and fat tissues originally described by Forbes (10).

To simulate RYGB surgery and intensive calorie restriction, P(t) was assumed to be single parameter value for t>0. To simulate the escalating doses of semaglutide and tirzepatide, P(t) = P_0_ + (P_max_-P_0_)t/T until t=T, where P_0_ is the initial drug effect and time t=T is the end of the dose escalation period, after which P(t) = P_max_ indicating the maximum drug effect. Predicted changes in energy intake and total energy expenditure were compared with mean data at corresponding time points, where available.

## Results

To simulate the calorie restriction intervention in the CALERIE phase 2 study (7), the best fit model parameters were k = 82 kcal/d per kg and P(t) = 830 kcal/d resulting in a body weight change trajectory that plateaued after about 12 months with simulated body fat being slightly greater than the observations (Figure 1A). Figure 1B shows a substantial initial decline of energy intake at the onset of the intervention followed by an exponential increased over time that closely matched observed intake change data obtained using the intake balance method (11). Simulated total energy expenditure rapidly decreased after the intervention onset and thereafter maintained an approximately constant level over the remainder of the 24-month intervention and matched the doubly labeled water energy expenditure data reasonably well.

**Figure 1.**
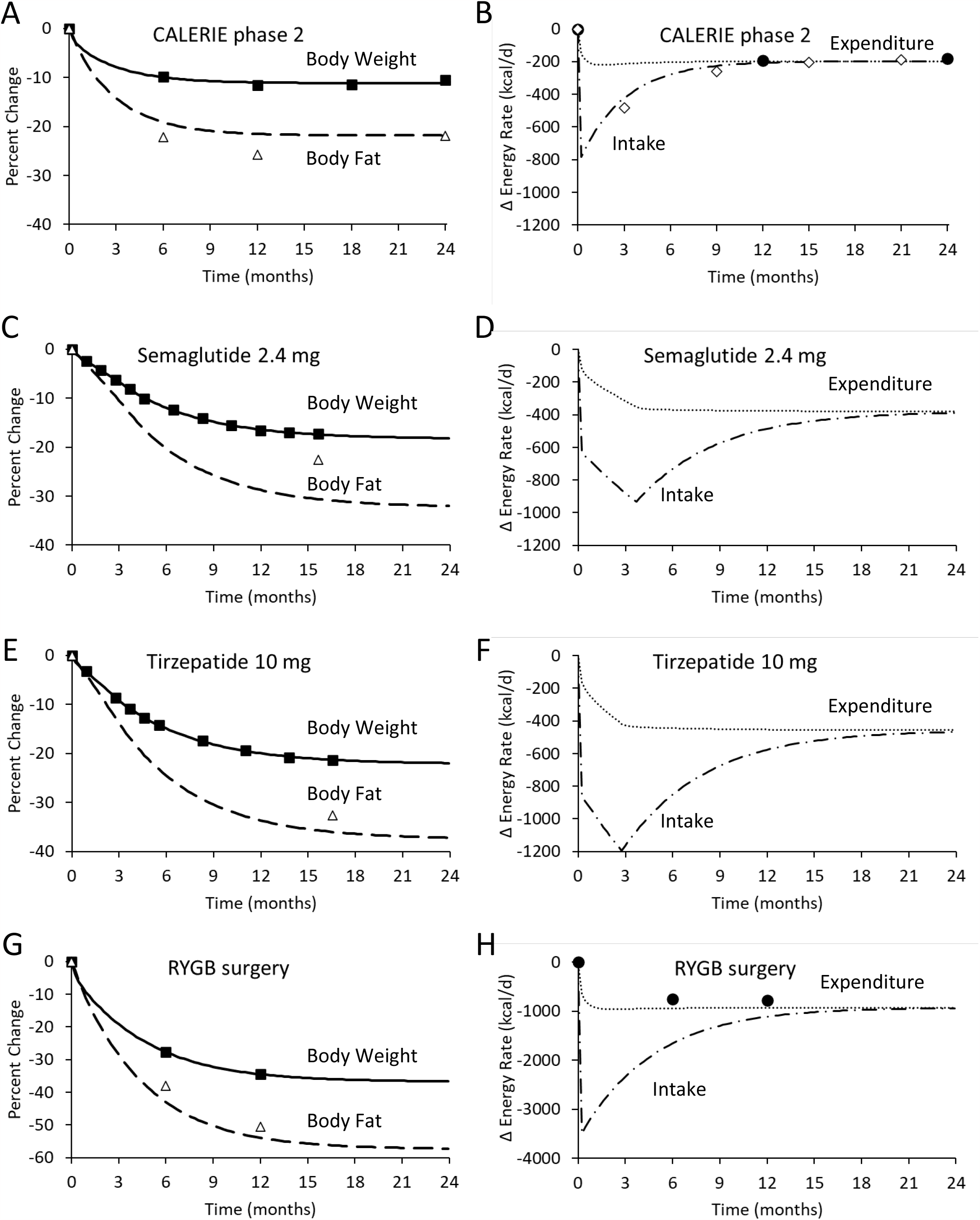
Body composition and energy balance dynamics during intensive calorie restriction (A and B), semaglutide 2.4 mg (C and D), tirzepatide (E and F), and RYGB surgery (G and H). Simulated body weight is depicted in the solid curves and was fit to the mean data depicted as closed boxes. The dashed curves are the simulated body fat changes and the DXA determined mean body fat changes are indicated by open triangles. Dotted curves indicate simulated energy expenditure changes and mean energy expenditure data are depicted by solid circles, where available. Dashed dotted curves represent the simulated energy intake changes and the open diamonds depict the mean energy intake data, where available.

Simulation of semaglutide therapy (2) was achieved using best fit parameters k = 49 kcal/d per kg, P_0_ = 610 kcal/d, and P_max_ = 1300 kcal/d after the 16-week dose escalation period resulting in an expected weight change plateau about 24 months after the intervention onset (Figure 1C). Unlike the calorie restriction simulation, the model predicted much lower levels of body fat than were observed. Energy intake decreased at the onset of the intervention and further declined during the dose escalation period after which intake exponentially increased until approaching the simulated energy expenditure towards the end of the 24-month simulation (Figure 1D).

Tirzepatide therapy was simulated using best fit parameters k = 48 kcal/d per kg, P_0_ = 830 kcal/d, and P_max_ = 1560 kcal/d after the 12-week dose escalation period (1) resulting in an expected weight change plateau after about 24 months (Figure 1E). Unlike the semaglutide simulation, tirzepatide therapy resulted in body fat measurements that were only slightly higher than the simulated values. Tirzepatide simulations resulted in a rapid initial decrease in energy intake that and further declined during the dose escalation period after which intake exponentially increased until approaching the simulated energy expenditure towards the end of the 24-month simulation (Figure 1F).

RYGB surgery was simulated using best fit parameters k = 58 kcal/d per kg and P = 3600 kcal/d resulting in the greatest weight losses of the simulated interventions and plateauing at 24 months with measured body fat changes were only slightly greater than the simulated values (Figure 1G) (8). Energy intake decreased by a very large amount after the surgery corresponding to almost no food intake immediately after the surgery followed by an exponential increase over time to approach total energy expenditure after about 24 months which decreased by ∼150 kcal/d more than was measured using a respiratory chamber.

## Discussion

The magnitude of the persistent intervention shifting the system from its baseline equilibrium following RYGB surgery was more than 4-fold greater than intensive calorie restriction and about double that of tirzepatide which was slightly larger than semaglutide. While intensive calorie restriction achieved the smallest intervention magnitude, the study participants exerted a substantial persistent effort to cut ∼800 kcal/d from their baseline diet. The exponential rise in energy intake after the start of the intervention shows that the same amount of effort to cut calories was met with increasing resistance as ongoing weight loss increasingly activated the feedback control circuit stimulating appetite. Eventually, increased appetite matched the persistent effort to cut calories and weight loss plateaued within 12 months.

Why did the other interventions result in prolonged weight loss periods with plateaus well after 12 months? Interestingly, the mathematical model suggests that the time to reach a weight plateau has nothing to do with the intervention magnitude P. Rather, a linearized version of the mathematical model used in the present study shows that the characteristic time scale of the system is given by *τ* = *ρ*/(*k* + *ε*), where ρ is the effective energy density of the weight change, ε defines the change in energy expenditure per unit weight change incorporating metabolic adaptation to weight loss, and k is the appetite feedback gain parameter (4). Decreasing k corresponds to a weakened appetite feedback control and results in a longer time to approach the weight plateau.

The preliminary estimate of k = 95 kcal/d per kg weight loss (5) was reasonably close to the best fit value of k = 83 kcal/d per kg weight loss achieved by calorie restriction in the CALERIE phase 2 trial and resulted in the weight plateau within 12 months. However, RYGB surgery, semaglutide, and tirzepatide all resulted in much lower values of k of ∼50-60 kcal/d per kg suggesting that these interventions weakened the feedback control of appetite and thereby resulted in a prolonged period of weight loss prior to the plateau.

Another way to increase the characteristic time τ to approach a weight plateau is for energy expenditure to decrease per unit weight loss to a lesser extent than expected corresponding to a lower-than-expected value of the parameter ε. However, because baseline ε is only ∼25 kcal/d per kg there is a limited ability to decrease ε. Thus, the observed prolongation of time to approach a weight plateau with RYGB surgery, tirzepatide, and semaglutide is most likely due to a weakening of the feedback control of appetite rather than an alteration of energy expenditure.

The simulated body fat changes generally matched the observations quite well except for the semaglutide intervention where body fat loss was substantially less than predicted for the observed weight loss. Thus, a greater than expected proportion of fat-free mass was lost with semaglutide treatment. More research is needed to investigate body composition changes during semaglutide treatment and whether the composition of fat-free mass lost has functional implications.

Simulated free-living energy intake and expenditure changes with calorie restriction matched the observations reasonably well, but RYGB surgery resulted in decreased expenditure measured using respiratory chambers that were slightly less than predicted by the model. This may be indicative of an effect of bariatric surgery to preserve energy expenditure or perhaps the respiratory chamber data did not fully capture the changes in free-living energy expenditure where physical activity is typically higher. Model simulations of energy intake and expenditure for tirzepatide or semaglutide treatment could not be compared with data because such measurements are not yet available.

In conclusion, varying model parameters P and k that influence energy intake allowed simulated body weight to fit body weight trajectories during a variety of weight loss interventions. The simulations indicated that RYGB surgery, tirzepatide, and semaglutide interventions weakened the feedback control of appetite unlike calorie restriction. The persistent magnitudes of these interventions were quantified and varied by more than 4-fold, but even the intensive calorie restriction intervention corresponded to a persistent effect to cut energy intake by ∼800 kcal/d over the two-year simulations. Interestingly, the model simulations do not predict any weight regain when assuming fixed values of the model parameters during the latter stages of the interventions. This suggests that explaining weight regain requires additional assumptions beyond the scope of the present study.

